# Deregulation of epigenetic factor, KDM6A, is involved during Candida albicans mediated delayed wound healing

**DOI:** 10.1101/2024.12.09.627481

**Authors:** Ankita Ghoshal, Awantika Shah, Kithiganahalli Narayanaswamy Balaji

**Author notes:** **Corresponding author:** Kithiganahalli Narayanaswamy Balaji, Department of Microbiology and Cell Biology, Indian Institute of Science, Bangalore – 560012, Karnataka, India. **Conflicts of Interest:** The authors declare no conflict of interest.

## Abstract

Deregulation of wound healing process leads to chronic non-healing wounds. Opportunistic fungal pathogens like *Candida albicans* is known to colonize open wounds leading to delayed, insufficient, or disorganized healing. In our study, we investigate the regulatory mechanisms orchestrating this *C. albicans* mediated delay of wound healing and underscore a critical role for the transcription factor c-AMP Response Element Binding Protein (CREB) in regulating pro-inflammatory cytokines and genes implicated in wound healing. Parallelly, we found the epigenetic factor, KDM6A (Ubiquitously Transcribed tetratricopeptide repeat on chromosome X, also known as UTX) to be downregulated upon infection and modulating expression of metalloproteinases during *C. albicans* governed wound healing impairment. Mechanistically, we observed enhanced recruitment of EZH2, a methyl transferase, over the promoters of KDM6A during infection, hence, governing its expression. Our findings, therefore, underscore a role of KDM6A and CREB in *C. albicans* mediated delayed wound healing.

## Introduction

Many of the protective defence mechanisms found in intact skin are lost in open skin wounds, leaving them vulnerable to microbial contamination^1^. Microbes generally present on the skin aid homeostasis, however, a commensal overgrowth or pathogen infiltration becomes detrimental to the natural process of wound healing. A fine regulation of genes implicated during tissue repair is necessary for effective wound healing^2^. During chronic wounds, elevated levels of Matrix Metallo Proteases (MMPs) with a concomitant decrease in the expression of their antagonists, tissue inhibitors of metalloproteases (TIMPs), have been reported^3^. Growth factors also play an important role^2^ and secretion of these factors by macrophages is very crucial during the early phase of wound repair^4^. The growth factors recruit fibroblasts and epithelial cells to the wound site^5^ ensuring tissue remodelling and subsequent healing.

A rise in pathogenic *Candida sp.* contamination of wounds is being reported^6^ and fungi were found to be present in approximately 23% of chronic wounds^7^ and in 80% of the diabetic foot ulcers. *Candida albicans* accounts for one of the most prevalent opportunistic fungal communities associated with fungus mediated pathogenicity in immunocompromised individuals^8^ and is known to cause open wound colonisation, oropharyngeal candidiasis, and vaginal candidiasis^9,10^. Further, emergence of antifungal drug resistance has prompted a new line of research into the role of fungus in delayed wound healing, as well as the potential host-directed mechanisms involved^11^. In a previous report published from our group, it has been shown that the regulation of certain genes that affect wound healing are perturbed at transcript levels upon *C. albicans* infection^12^ leading to delayed wound healing. We were interested to further investigate the possible mechanisms that could be orchestrating *C. albicans* mediated delayed wound healing.

In the recent times, epigenetic regulators and transcription factors have received a lot of attention in the context of diseases and infections^13^ with histone modifications play a significant role in determining gene expression^14^. KDM6A (ubiquitously transcribed tetratricopeptide repeat on chromosome X, also known as UTX), a histone demethylase that removes the di- and tri-methyl groups from histone H3K27 residue has been shown to be upregulated in normal homeostatic wound healing^15^. However, the functional implications of KDM6A on wound healing have not been explored.

Various transcription factors are responsible for maintaining homeostasis in cells, and they have been also reported to play a major role in wound healing^16^. One of the major transcription factors that plays a central role in regulating important biological functions is CREB (cAMP response element binding protein)^17^. CREB, a master transcription regulator, plays numerous roles in immune system function^17^. CREB, though widely known for its role cell proliferation, differentiation, and survival, has also been reported to regulate several immunological responses namely activation of pro-inflammatory mediators upon external insult, macrophage polarization, T cell responses^17^. Pro-inflammatory cytokines are reported to be persistently upregulated upon *C. albicans* infection. We were therefore interested in understanding whether CREB might be playing an immunomodulatory role in *C. albicans*-mediated delayed wound healing.

In our study, we report that the downregulation of H3K27me3 demethylase KDM6A upon *C. albicans* infection is primarily responsible for the repressed expression of growth factors that are essential for homeostatic wound healing with deposition of enhanced repressive mark H3K27me3 on the promoters of these genes upon infection. Parallelly, we also report a significant role of CREB activation in modulating the enhanced expression of pro-inflammatory cytokines upon *C. albicans* infection, which contributes significantly to delayed healing.

## Results

### Deregulation of inflammatory signature during *C. albicans* mediated delayed wound healing

Punch biopsy model was utilized for studying the process of wound healing in mice^18^. Briefly, two symmetrical circular excisional wounds of comparable thickness were created on shaved dorsal skin of the mouse, after which one of the wounds was infected with *C. albicans* and the other wound was administered PBS as control. Upon careful monitoring of the wounds, we observed that wounds that were administered PBS underwent fast healing whereas wounds infected with *Candida* displayed markedly slow healing (**Fig. 1A**). Persistent expression of pro-inflammatory cytokines coupled with repressed expression of growth factors is one of the hallmarks of chronic wounds^19^. Downregulation of anti-inflammatory cytokines in *C. albicans* infected wounds^20^, usually associated with wound resolution, implicate their role in delaying wound healing process (**Fig. 1B**). We also observed reduced expression of genes implicated in promoting wound healing, mainly growth factors and TIMPs, in infected wound samples as compared to uninfected wounds (**Fig. 1C**). The effect of delayed wound healing was prominent despite the upregulation of pro-inflammatory factors like *Ccl2, Tnfa, Il1b* and *Nos2* in the *C. albicans* infected wound.

**Fig. 1:**
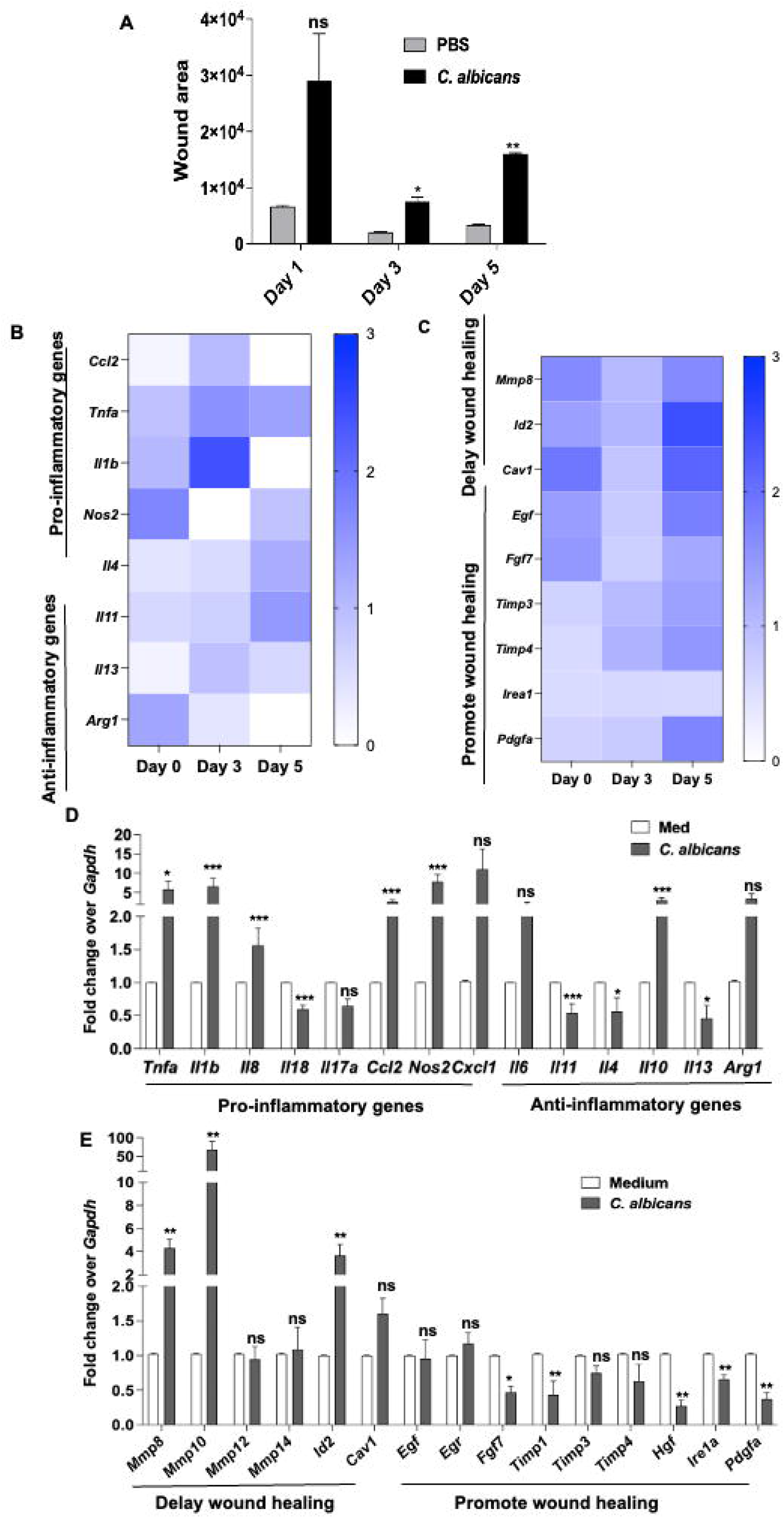
*C. albicans* leads to delay in wound healing with aberrant expression of inflammatory cytokines and genes implicated in wound healing. (A) Photographs of wounds were analyzed using ImageJ software on indicated days to determine area of wounds treated with PBS or those infected with *C. albicans*. (B, C) RNA was isolated from wound samples and qPCR analysis was done for pro-inflammatory and anti-inflammatory chemokines and wound healing markers. (D,E) Mouse peritoneal macrophages were infected with *C. albicans* for 12h and levels of pro and anti-inflammatory chemokines and wound healing genes was assessed. Data represent the mean ± SEM of at least two independent experiments having at least three mice in each group. ns, non-significant, *p < 0.05, **p < 0.005, ***p<0.0001, Student’s t-test.

Since macrophages are known to play an important role during wound healing process^20^, we sought to assess the levels of genes implicated in the process in an *ex vivo* model. In this regard, macrophages were infected with *C. albicans* and in corroboration with our *in vivo* data, we observed an increase in the expression of *Mmp8*^21^, known to degrade collagen and *Mmp10*, both previously associated with aberrant wound healing. Expression of pro-inflammatory genes were significantly pregulated in infected macrophages as compared to uninfected macrophages (**Fig. 1D**). This was accompanied by a marked reduction in the levels of TIMPs (*Timp1, Timp3, Timp4*) and growth factors (*Hgf, Pdgfa, Egf* & *Fgf7*) known to accelerate the process of wound healing (**Fig. 1E**). Hence, a heightened inflammatory response, dysregulation of balance between the expression of MMPs and TIMPs, and loss of growth factors are associated with delayed wound healing during *C. albicans* infection.

### CREB activation upon *C. albicans* infection regulates genes implicated in wound healing

With the aim to dissect molecular players that might be responsible for the observed modulation of gene expression leading to wound healing delay, we investigated the role of CREB, a master transcriptional regulator, during *C. albicans* infection. Levels of p-CREB were significantly upregulated post *C. albicans* infection both *in vivo* and *ex vivo* (**Fig. 2A, B**). CREB, in its active phosphorylated form (on serine-133), binds to the promoters of its target genes and activates the transcription status of those genes^17^. Therefore, we wanted to assess whether CREB could be regulating the levels of genes observed to be modulated during *C. albicans* mediated delayed wound healing process. We observed that increased expression of genes implicated in delayed wound repair post *C. albicans* infection was abrogated upon CREB activity inhibitor pre-treatment in macrophages (**Fig. 2C, D**). Application of CREB activity inhibitor to infected wounds led to their faster healing as compared to untreated infected wounds (**Fig. 2E**). H&E staining suggested that in *C. albicans* infected wound, re-epithelialization is scant, tissue fibrosis is high coupled with scant levels of granulation tissue deposition whereas the inhibitor treated wounds have a fast re-epithelialization rate, abundant granulation tissue deposition and have mostly regained their normal skin texture (**Fig. 2F**). Hence, CREB activity delays wound healing process during *C. albicans* infection.

**Fig. 2:**
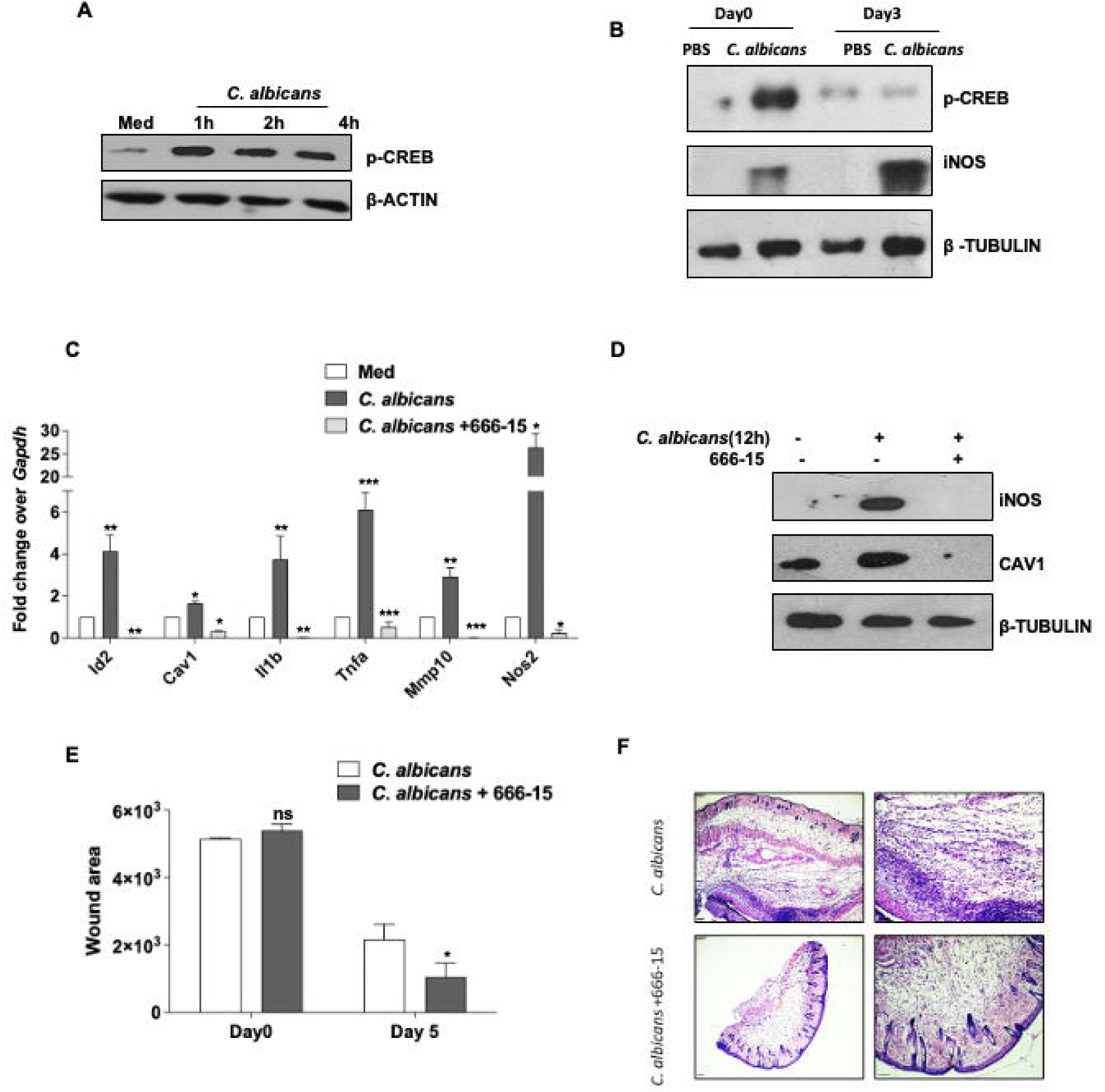
p-CREB regulates *C. albicans* mediated delayed wound healing: (A) Mouse peritoneal macrophages were infected with *C. albicans* and phosphorylation status of CREB was assessed at 1h, 2h and 4h post infection. (B) Levels of p-CREB and iNOS in infected wounds were assessed. Mouse peritoneal macrophages were pre-treated with CREB activity inhibitor 666-15(5μM) and then infected with *C. albicans* for 12h infection and (C) levels of indicated genes were assessed at transcript levels and (D) protein levels. (E) Infected wounds were treated with p-CREB inhibitor 666-15 (20mM) on alternate days post wounding and assessed. Quantification of wound area (F) H&E staining of infected wounds with and without CREB inhibitor treatment is shown. Data is representative of least three mice in each group. All *ex vivo* data represent the mean ± SEM of least three biological replicates. All *ex vivo C. albicans* infections were performed at MOI of 1:1 (macrophage/yeast ratio). *p < 0.05, **p < 0.005, ***p < 0.0001; one-way ANOVA. ns, non-significant, *p<0.05, Student’s t-test.

### Cannabinoid receptor 2 (CB2) signalling regulates p-CREB in macrophages during *C. albicans* infection

We next wanted to assess the mechanism by which p-CREB levels might be regulated by *C. albicans* infection in macrophages. Reports suggest a role for cannabinoid receptor 2 (CB2) signalling in activating p-CREB^22^.The endocannabinoids like 2-AG (2-arachedonyl glycerol), the primary CB2 agonist, has been reported to act on various immune cell types by modulating leukocyte recruitment functions such as chemokine release, fibronectin adhesion, and migration. 2-AG’s has also been reported to activate pro-inflammatory response in cells^23^.

We found that upregulation of p-CREB levels upon *C. albicans* infection was compromised upon pre-treatment of cells with CB2 inhibitor SR-144528 (**Fig. 3A**). Further, p-CREB levels were also upregulated in macrophages treated with CB2 agonist JWH-133 (**Fig. 3B**). Upon silencing of CB2 in macrophages, we found that levels of iNOS and CAV1, two key proteins that have been implicated in delayed wound healing^24,25^, were downregulated post *C. albicans* infection (**Fig. 3C**). Interestingly, CB2 receptor levels remained unchanged post infection (**Fig. 3D &E**). Hence, Cannabinoid Receptor 2 mediated signalling regulates CREB activation upon *C. albicans* infection.

**Fig. 3:**
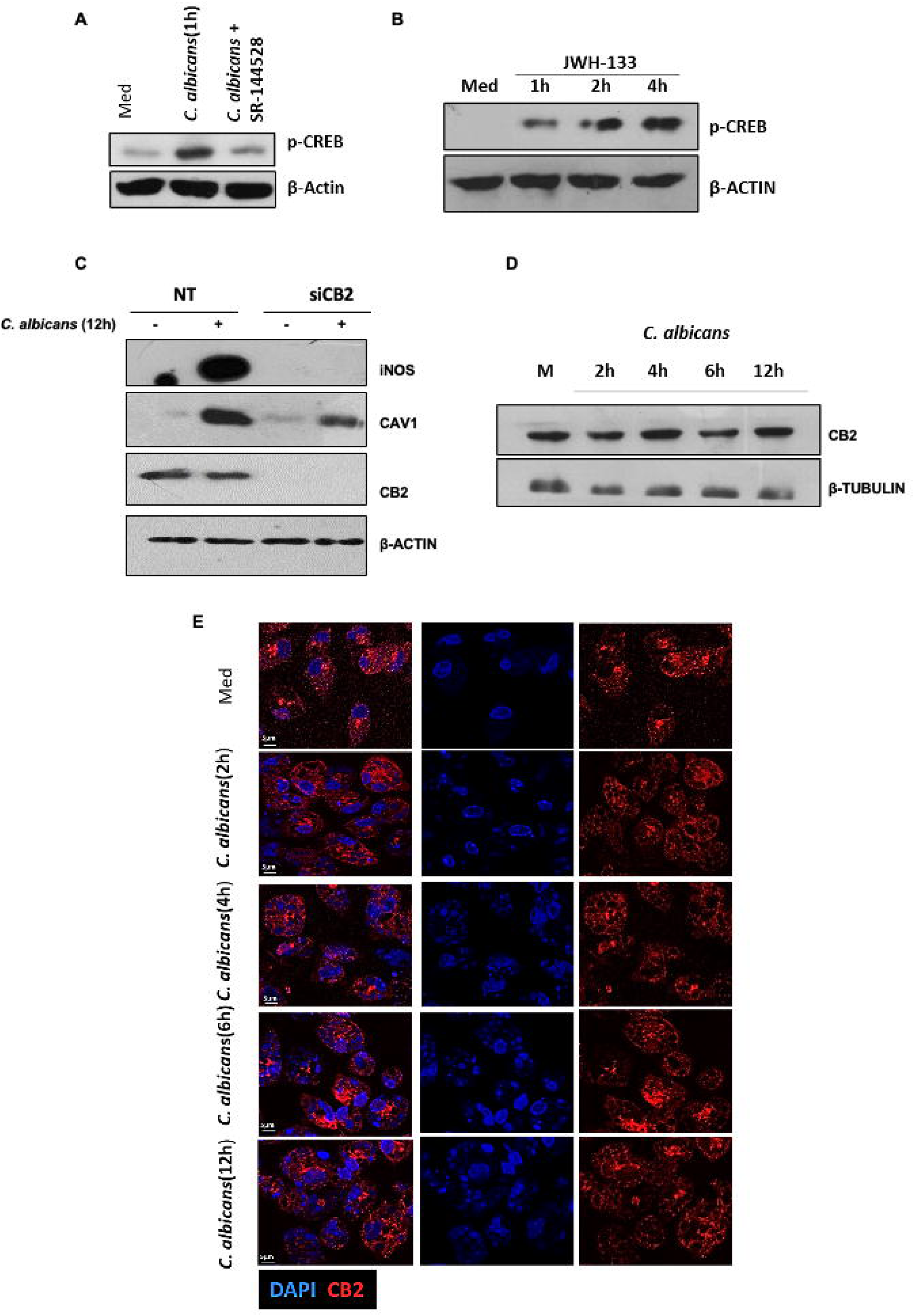
p-CREB is regulated by Cannabinoid receptor 2 mediated signaling: (A) Mouse peritoneal macrophages were pre-treated with CB2 inhibitor SR-144528 (10μM). Following 2h of infection with *C. albicans* post inhibitor pre-treatment, p-CREB levels were assessed. (B) Mouse peritoneal macrophages were treated with CB2 receptor specific agonist JWH-133 (10μM) for 1h, 2h and 4h respectively, post which p-CREB protein levels were assessed. (C) Mouse peritoneal macrophages were transfected with CB2 siRNA. Following 12h of infection with *C. albicans* post inhibitor pre-treatment, iNOS and CAV1 protein levels were assessed. Data represent the mean ± SEM of two biological replicates (D&E) Mouse peritoneal macrophages were infected with *C. albicans* for 2h, 4h, 6h and 12h respectively, post which CB2 protein levels were assessed. All *ex vivo* data represent the mean ± SEM of at least three biological replicates unless mentioned otherwise. All *ex vivo C. albicans* infections were performed at MOI of 1:1 (macrophage/yeast ratio).

### EZH2 regulates KDM6A levels during *C. albicans* mediated delayed wound healing

We observed the upregulation of pro-inflammatory cytokines upon *C. albicans* infection in both *ex vivo* and *in vivo* scenario. Usually, upregulation of pro-inflammatory factors is followed by upregulation in growth factors like *Fgf7*, *Pdgfa, Egf*, which tackles the pro-inflammatory state of the wound and drives it towards homeostasis and normal healing^5^. Interestingly, we observed downregulation in the expression levels of growth factors *Pdgfa*, *Egf* and *Fgf7* post *C. albicans* infection (**Fig. 1C**). Previous study from our lab has shown that EZH2, a histone methyl transferase, that belongs to the Polycomb Repressive 2 Family Class of genes deposits the repressive H3K27me3 mark on histones along with enhanced recruitment on the promoters of genes which promote wound healing upon *C. albicans* infection in macrophages^12^. The increased recruitment of EZH2 on the promoters of these genes leads to their repression, which might be one of the factors majorly responsible for delayed wound healing under these conditions. Homeostasis between deposition of these repressive marks by a histone methyltransferase and simultaneous removal of those marks by a histone demethylase is critical for proper gene expression^26^. In this regard, reports suggest EZH2 regulates a histone demethylase, KDM6A, to govern various gene expression changes^27^.

We first evaluated the deposition of repressive H3K27me3 mark on the promoter of KDM6A and found enhanced deposition of the repressive mark upon *C. albicans* infection (**Fig. 4A**). Enhanced deposition of H3K27me3 transferase EZH2 (Enhancer of Zeste Homolog 2) was also observed on KDM6A promoter post *C. albicans* infection in macrophages (**Fig. 4B**). To assess the regulation of KDM6A, we utilized GSK126 (EZH2 inhibitor) in macrophages and observed that the downregulated levels of KDM6A observed upon *C. albicans* infection was significantly rescued in the inhibitor pre-treated macrophages (**Fig. 4C, E, F**). Since EZH2 inhibition lowers the global levels of H3K27me3 mark, the levels of the same were also assessed as a positive control to determine inhibitor efficiency (**Fig. 4D**). The levels of IRE1A, a wound healing promoting marker that was earlier shown, were evaluated as well with respect to *C. albicans* infection upon GSK126 (EZH2 inhibitor) pre-treatment in macrophages. IRE1A was chosen as a positive control because it is also a reported downstream target of KDM6A ^28^. The downregulated levels of IRE1A upon *C. albicans* infection was significantly recovered in inhibitor pre-treated cells (**Fig. 4 G, H**). Hence, EZH2 regulates KDM6A downregulation upon *C. albicans* infection.

**Fig. 4:**
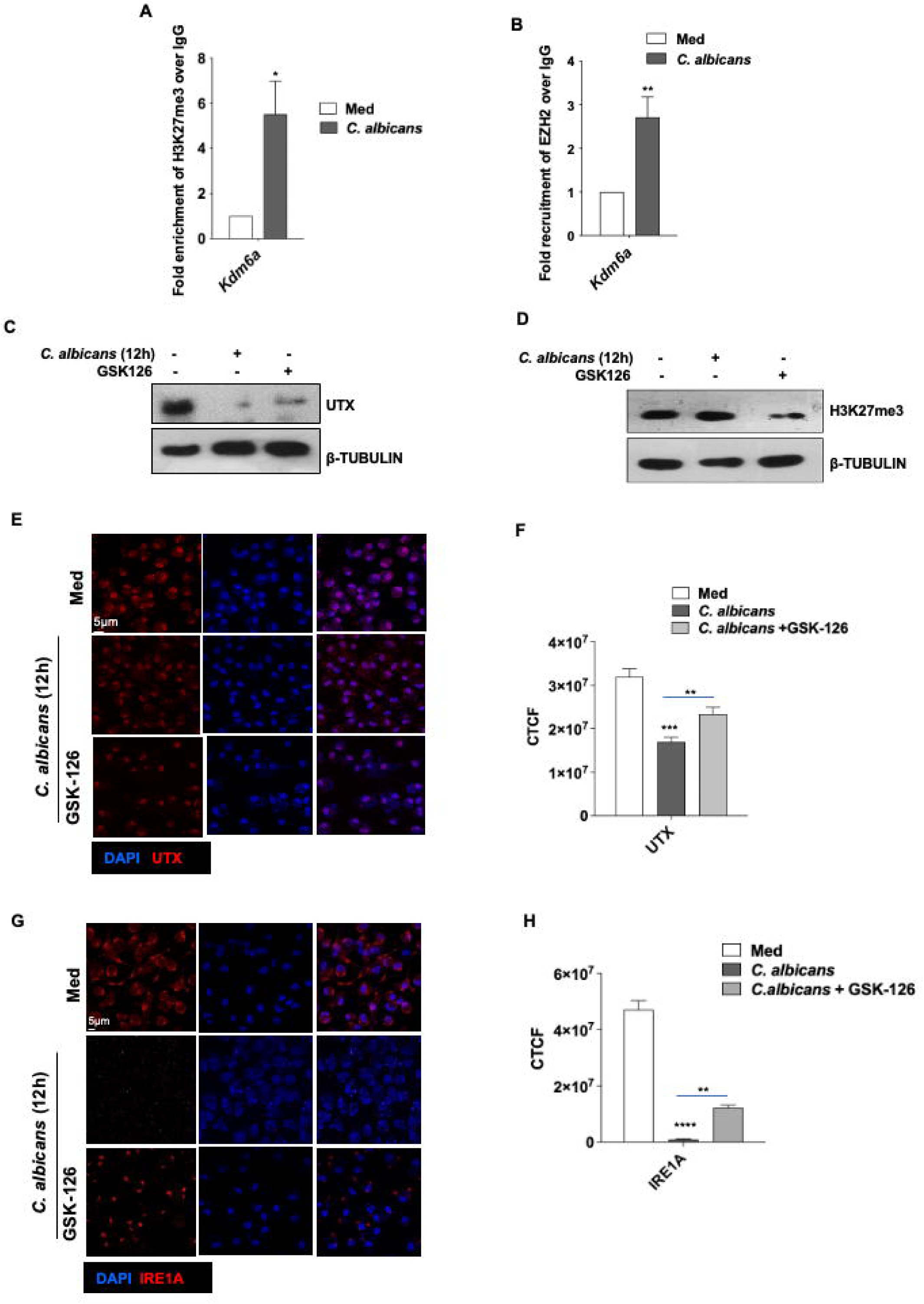
Histone methyltransferase EZH2 regulates *Kdm6a* levels upon *C. albicans* infection: (A-B) Mouse peritoneal macrophages were infected with *C. albicans*. Following 12h of infection, (A) H3K27me3 enrichment and (B) EZH2 recruitment was assessed on the promoter of *Kdm6a* by chromatin immunoprecipitation. (C) Mouse peritoneal macrophages were pre-treated with EZH2 inhibitor GSK126(10μM). Following 12h of infection with *C. albicans* after inhibitor pre-treatment, UTX and (D) H3K27me3 levels were assessed. (E) Mouse peritoneal macrophages were pre-treated with GSK126(10μM). Following 12h of infection with *C. albicans* after inhibitor pre-treatment, UTX protein levels were assessed by immunostaining. (F) quantification of the same data. (G) Mouse peritoneal macrophages were pre-treated with GSK126(10μM). Following 12h of infection with *C. albicans* after inhibitor pre-treatment, IRE1A protein levels were assessed (H) quantification of the same data. All *ex vivo* data represent the mean ± SEM of at least three biological replicates unless mentioned otherwise. All *ex vivo C. albicans* infections were performed at MOI of 1:1 (macrophage/yeast ratio). *p < 0.05, **p < 0.005, ***p < 0.0001, *p < 0.05. Student’s t-test for A and B, ONE WAY ANOVA for F and H.

### Deregulated expression of histone demethylase KDM6A perturbs homeostatic regulation in macrophages during *C. albicans* infection

Next, to understand the implication of KDM6A in wound healing process, we assessed for change in expression of KDM6A upon *C. albicans* infection and found it to be significantly downregulated in infected wounds as opposed to uninfected wounds (**Fig. 5A**). Moreover, KDM6A levels were also downregulated upon *C. albicans* infection in macrophages at both transcript (**Fig. 5B**) and protein levels (**Fig. 5C-E**).

**Fig. 5:**
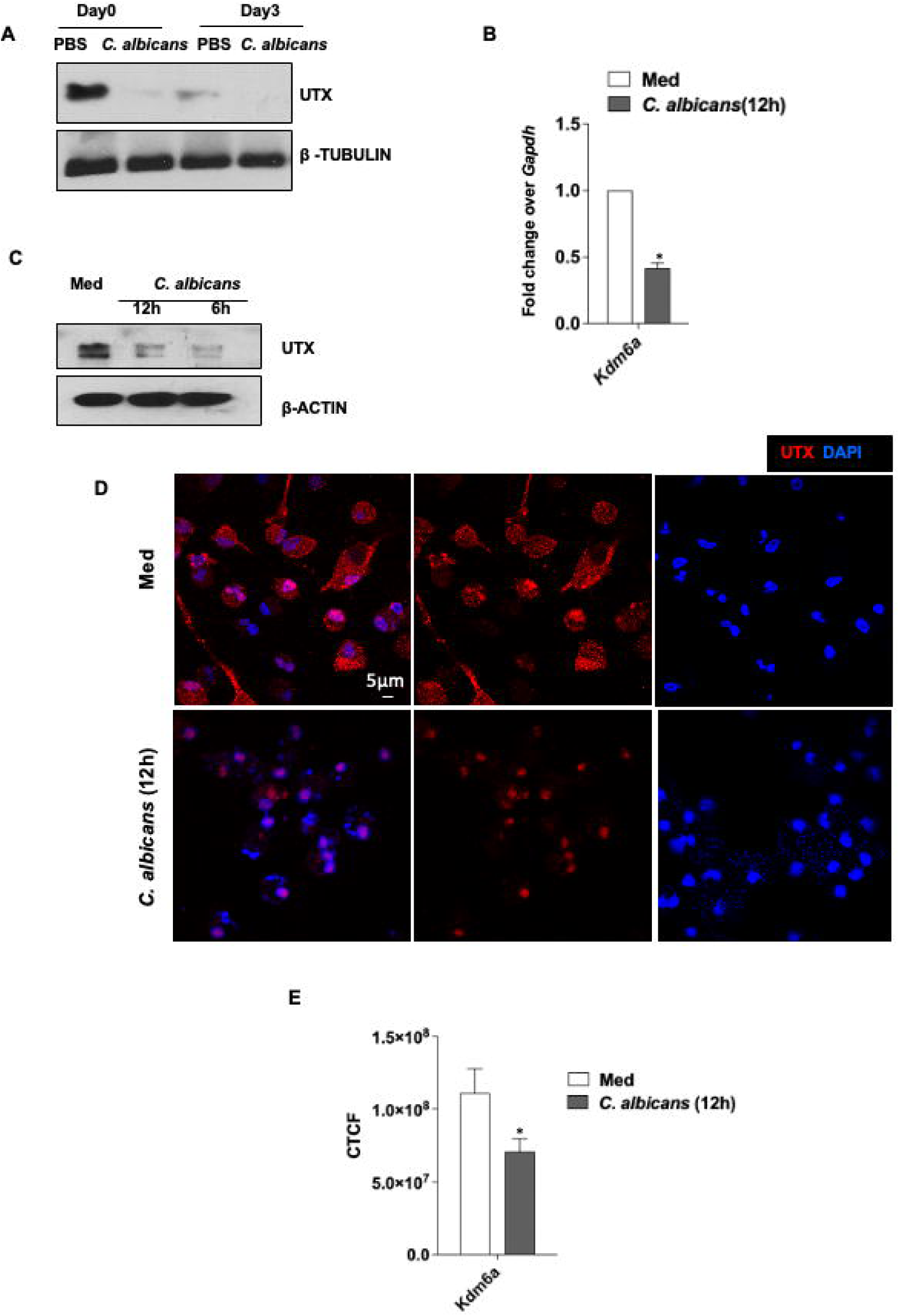
Histone demethylase *Kdm6a* is downregulated upon *C. albicans* infection: (A) KDM6A levels were assessed at protein levels in wound samples. (B) Mouse peritoneal macrophages were infected with *C. albicans* for 12h and KDM6A levels were assessed at transcript levels and protein levels by (C) immunoblotting and (D) immunostaining. (E) Quantification. All *ex vivo* data represent the mean ± SEM of at least three biological replicates. All *ex vivo C. albicans* infections were performed at MOI of 1:1 (macrophage/yeast ratio). *p < 0.05, Student’s t-test.

To explore a regulatory relationship, we assessed whether KDM6A overexpression would be able to rescue the levels of genes that were repressed upon *C. albicans* infection and critical for tissue repair. For this, we utilized KDM6A overexpression plasmid, and observed that the expression levels of the genes implicated in wound healing process that were downregulated upon *C. albicans* infection were rescued in the KDM6A overexpressing cells (**Fig. 6A-C**). We further evaluated the effect of KDM6A overexpression on *C. albicans* survival in macrophages and *C. albicans* burden in macrophages was significantly decreased upon KDM6A overexpression in macrophages (**Fig. 6D**). Parallelly, we assessed the recruitment of KDM6A on promoters of genes promoting wound healing *(Pdgfa, Egf, Fgf7, Ire1a*) and found significantly reduced occupancy of KDM6A upon *C. albicans* infection in macrophages (**Fig. 6E**) coupled with an enhanced occupancy of repressive H3K27me3 mark (**Fig. 6F**). Therefore deregulated KDM6A expression during *C. albicans* infection affects genes implicated in wound healing.

**Fig. 6:**
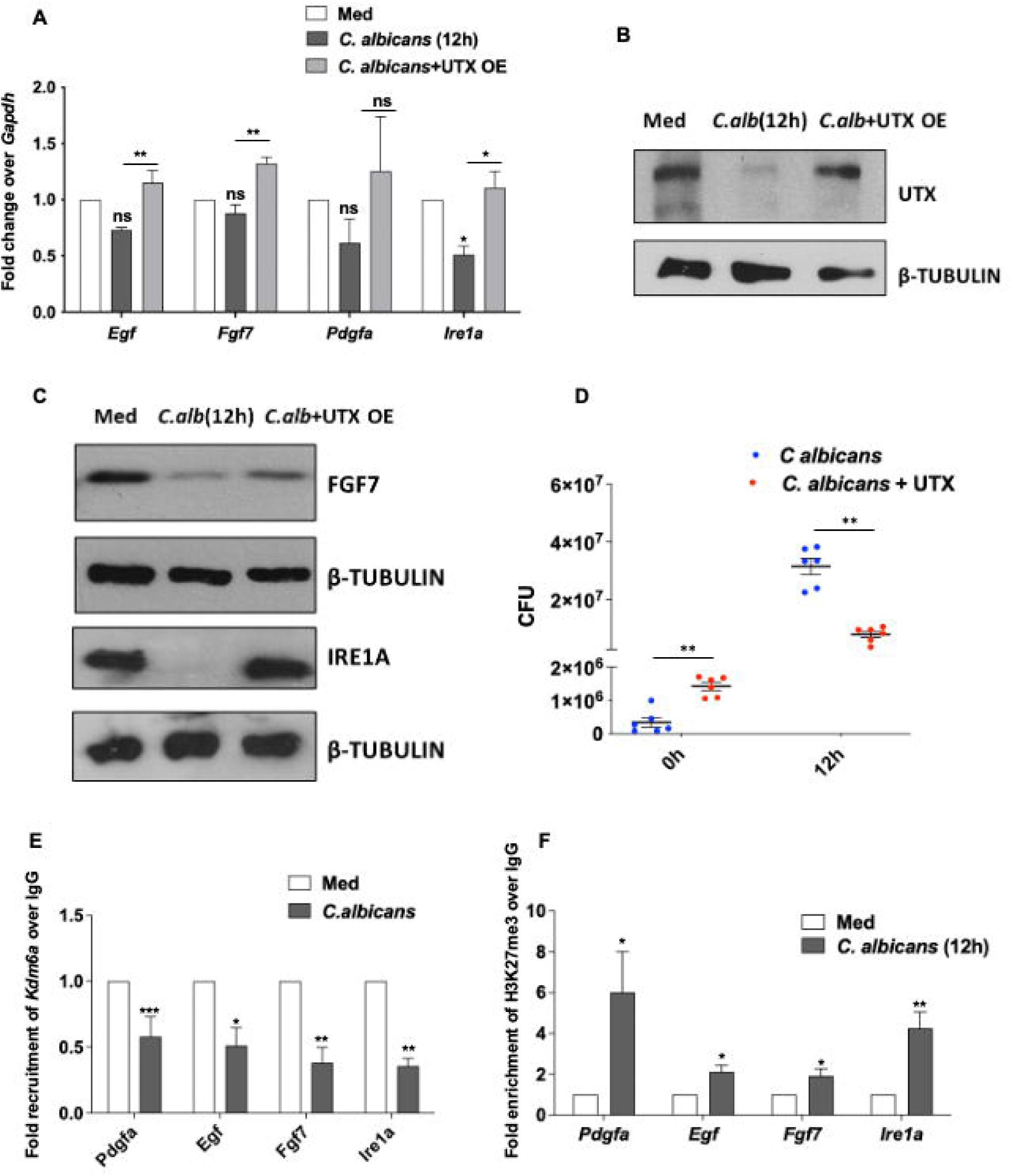
KDM6A downregulation deregulates wound healing promoting genes in macrophages: (A-C) RAW264.7 were transfected with vector control (Med) and *Kdm6a* over-expression construct prior to infection with *C. albicans*. Following 12h of infection, levels of indicated genes were assessed at transcript and protein levels respectively. Empty vector transfected cells were used as negative control. (D) RAW 264.7 cells were transfected with vector control and KDM6A overexpression construct and intracellular survival of *C. albicans* was assessed at 0h and 12h post infection. (E-F) Mouse peritoneal macrophages were infected with *C. albicans*. Following 12h of infection, (E) recruitment of KDM6A and (F) H3K27me3 repressive mark were assessed on the promoter of indicated genes by means of chromatin immunoprecipitation. All *ex vivo* data represent the mean ± SEM of at least three biological replicates. All *ex vivo C. albicans* infections were performed at MOI of 1:1 (macrophage/yeast ratio). ns, non-significant, *p < 0.05, **p < 0.005, ***p < 0.0001, Student’s t-test.

## Discussion

In our study, we focused on understanding the possible mechanisms involved in *C. albicans*-mediated delayed wound healing. There are four phases of healing, namely homeostasis, inflammation, re-epithelialization, and tissue remodeling^29^. *Pdgf*, *Fgf7*, *Egf,* and other growth factors are secreted after wounding, causing monocytes, macrophages, and inflammatory cells to be recruited to the wound^5^. The inflammatory phase of wound healing is critical in determining whether a wound will heal or enter a chronic non-healing condition^30^. Malnutrition, cardiovascular disease, obesity, microbial colonization of the wound, diabetes, autoimmune illnesses, maceration, etc. are usually associated with defective or delayed wound healing^31^. Among these, microbial colonization of the wound plays an important role in the delayed healing process^32^. In our work, we observed that both *C. albicans* infected macrophages and wounds have elevated expression of genes known to inhibit homeostatic wound healing. This was also accompanied by increased expression of pro-inflammatory cytokines, resulting in a protracted inflammatory state that may be detrimental for scar tissue resolution. We set out to investigate the underlying causes of the elevated expression of these genes.

Reports suggest that transcription factors play an important role in determining the inflammatory landscape of wounds^16^. In host immune system, CREB, a master transcription regulator, has a wide array of effector actions and has been reported to modulate pro-inflammatory cytokines in cells^17,33^. In our study, we show that *C. albicans* infection increases p-CREB levels and inhibition of CREB transcriptional activity resulted in the downregulation of the aforementioned factors. Application of a CREB inhibitor to infected wounds resulted in faster wound healing. As a result, these findings indicate that CREB activation in response to *C. albicans* infection causes the wound to enter a protracted inflammatory stage. Next, we wanted to assess the signaling modulators that might facilitate p-CREB overexpression following *C. albicans* infection and found Cannabinoid receptor 2 mediated signaling to regulate CREB activity. CB2 mediated signaling has been extensively studied in the context of many diseases and infections^22,24^.

When pro-inflammatory cytokines are overexpressed in wound tissue, expression of growth factors are elevated to counteract their effects and reduce inflammation^19^. However, we found that the expression of these growth factors was suppressed in infected wounds and macrophages. Epigenetic changes are one of the key variables identified to regulate gene expression^34^. Multiple investigations have highlighted the crucial role that chromatin remodelers and histone modifications play in determining the expression of genes^35^. EZH2, a histone methyl transferase, has previously been demonstrated to suppress the expression of these genes in *C. albicans* infected macrophages^12^. EZH2 has been shown to transfer H3K27me3 marks on the promoters of numerous genes, leading them to be repressed. The alteration of histone lysine residues is an extremely dynamic process. The methylation marks mentioned above can be ‘erased’ by protein lysine demethylases^36^. Proper regulation of this “writer-eraser” activity is essential for a gene to express normally. To date, little is known about modulation of histone demethylases by *C. albicans* and its impact on delayed wound healing.

In our study, we show that *C. albicans* infection downregulates the expression of histone demethylase KDM6A in macrophages and infected wounds. This causes the repression of growth factors implicated in wound healing including *Pdgfa, Egf, and Fgf7*. We also show that EZH2 regulates KDM6A levels during *C. albicans* infection. EZH2 deposits elevated H3K27me3 marks on the promoter of *Kdm6a*, causing it to be downregulated. Hence, we showcase a role of histone demethylase KDM6A and transcription factor CREB in regulating *C. albicans* mediated delayed wound healing. This, in turn, opens up multiple avenues for future host directed therapeutic interventions against this condition.

## Methods

### Ethics statement

Animal experiments (mice) were conducted following approval from the IAEC (Institutional Ethics Committee for Animal Experimentation) and experiments using *Candida albicans* from the IBSC (Institutional Biosafety Committee). All the protocols for animal care and use were in adherence to the guidelines set by the CCSEA (formerly CPCSEA), (Committee for Control and Supervision of Experiments on Animals), Government of India.

### Cells, mice, and *C. albicans*

In this study, both male and female BALB/C mice were used. The mice were housed at the Central Animal Facility of Indian Institute of Science (IISc). Mice were given an intraperitoneal injection of 1 ml of 8% Brewer’s thioglycollate. After 4 days, mice were sacrificed, and peritoneal cells were harvested by lavage from the peritoneal cavity with ice-cold PBS. The cells were grown in DMEM (Gibco-Invitrogen/Thermo Fisher Scientific) with 10% FBS (Gibco-Invitrogen/Thermo Fisher Scientific) for 24 hours. Microbial Type Culture Collection, IMTECH, Chandigarh, India provided us with *C. albicans* (MTCC 4748) strain and grown in Yeast extract, Peptone and Dextrose (YPD) media at 30°C. *C. albicans* was counted for infection using a Neubauer chamber. All *C. albicans* infections *in vitro* were done at a MOI of 1:1 (macrophage/yeast ratio).

### Immunoblotting

Cells were washed with ice-cold PBS, scraped from the culture dish, and pelleted down by centrifugation. RIPA buffer (50 mM Tris-HCl, pH 7.4, containing 1 % NP-40, 0.25 % Sodium deoxycholate, 150 mM NaCl, 1 mM EDTA, 1 mM PMSF, 1 mg/ml aprotinin, 1 mg/ml leupeptin, 1 mg/ml pepstatin, 1 mM Na_3_VO_4_, and 1 mM NaF) was used to lyse cell. Lysed cells were centrifuged at 13,000 rpm for 15 minutes at 4°C to obtain whole cell lysate and protein was estimated using Bradford’s method of protein estimation (cells or tissue). SDS-PAGE electrophoresis was used to separate equal amounts of protein from each sample, which was then transferred to PVDF membranes (Millipore, USA) using the semidry (Bio-Rad, USA) method. Membranes were blocked in TBST [20 mM Tris-HCl (pH 7.4), 137 mM NaCl, and 0.1 % Tween 20] for 60 minutes with 5 % non-fat dry milk powder (bovine, Sigma-Aldrich). The blots were incubated overnight at 4°C with a diluted primary antibody in TBST containing 5% BSA. After washing with TBST, the blots were incubated for 4 hours with the appropriate secondary antibody conjugated to HRP (Jackson Immunoresearch USA) diluted in 5% nonfat milk. Following additional TBST washes, the immunoblots were developed with ECL reagent (Perkin Elmer, USA). β-ACTIN and β-TUBULIN were used as the loading control.

### Chromatin immunoprecipitation assay

The chromatin immunoprecipitation (ChIP) assays were performed according to a protocol modified by Upstate Biotechnology. Macrophages were fixed in 1.42 percent formaldehyde for 7 minutes at room temperature and then inactivated with 125 mM glycine. Cells were lysed in 0.1 percent SDS lysis buffer (containing 1 percent Triton X-100), and chromatin was sheared using a Bioruptor (Diagenode) Sonicator (high power, 70 cycles of 30 s of pulse on and 45 s of pulse off). Chromatin extracts containing DNA fragments averaging 500 bp in size were immunoprecipitated using specific antibodies or isotype IgG. Wash Buffer A (50 mM Tris-HCl [pH 8], 500 mM NaCl, 1 mM EDTA, 1 percent Triton X-100, 0.1 percent sodium deoxycholate, 0.1 percent SDS, and protease/phosphatase) and Wash Buffer B (50 mM Tris-HCl [pH 8], 1 mM EDTA, 250 mM LiCl, 0.5 percent SDS and 0.1 M NaHCO3) were used to wash immunoprecipitated complexes. Following RNAse A and Proteinase K treatment of the eluted samples, DNA was precipitated using the phenol/chloroform/ethanol method. Purified DNA was examined using quantitative real-time reverse transcription PCR. All test-sample values were normalised to the specific gene in input and IgG pull down and expressed as fold change.

### Immunofluorescence

Peritoneal macrophages were seeded on coverslips and infected for 12 hours with *C. albicans* at a MOI of 1:1 (macrophage/yeast ratio titrated for imaging purposes). Following that, the cells were fixed with 3.7 % formaldehyde. After blocking with 2% BSA, the coverslips were stained with specific Abs overnight at 4°C. For 2 hours, the coverslips were incubated with Alexa 555-conjugated secondary Ab, and the nuclei were stained with DAPI. Glycerol was used as the mounting medium for the coverslip. Confocal images were captured using a plan apochromat 63/1.4 oil differential interference contrast objective (ZEISS) on a Zeiss LSM 710 Meta confocal laser scanning microscope (ZEISS), and images were analyzed using ZEN 2009 software. Images were also captured using a plan apochromat 63/1.4 oil differential interference contrast objective on the Confocal Leica TCS SP5 microscope. We used a free hand selection tool to select the cells for analysis and measured the area-integrated intensity and mean grey value. The background values were calculated using the area around the cells that did not have fluorescence. The corrected total cell fluorescence (CTCF) was calculated as follows: CTCF stands for Integrated intensity-(area of selected cell X Mean fluorescence of background reading).

### Quantitative Real-Time RT-PCR

Macrophages were treated or infected as indicated and total RNA was isolated using TRI reagent (T9424, Sigma-Aldrich, USA). For RT-PCR, 1 μg of total RNA was converted into cDNA using First Strand cDNA synthesis kit (M3682, Promega). Quantitative real-time RT-PCR was performed with SYBR Green PCR mixture (F416, Thermo Fisher Scientific). *Gapdh* was used as internal control.

### Reagents and antibodies

All general chemicals and reagents were obtained from Sigma-Aldrich/ Merck Millipore, HiMedia, and Promega. Tissue culture plastic ware was obtained from Thermofischer Scientific, Jet Biofil or Tarsons India Pvt. Ltd., and Corning Inc. siRNAs against CB2 were obtained from Eurogentech. Sigma-Aldrich supplied HRP-tagged anti-ACTIN (A3854) and 4′,6-Diamidino-2-phenylindole dihydrochloride (DAPI). Cell Signaling Technology supplied anti-p-CREB, anti-UTX, anti-EZH2, and anti-H3K27me3 antibodies (USA). Abcam (USA) supplied the anti-CB2, anti β-TUBULIN, anti-IRE1A, and anti-FGF7 antibodies. Cloud-clone CORP provided anti-Caveolin1 antibody (USA). Jackson ImmunoResearch provided anti-rabbit IgG and anti-mouse IgG. (USA). Thermo Fisher Scientific provided Lipofectamine 3000. CREB inhibitor 666-15 was obtained from Merck Millipore (Cat. No.: 5383410001). CB2 inhibitor SR-144528 was obtained from Cayman Chemicals.

### Plasmids and constructs

#### KDM6A overexpression plasmid construct was obtained from Addgene

##### Transient transfection studies

Macrophages were transfected for 6 hours with Lipofectamine 3000/ PEI with the indicated constructs (UTX OE) or with 100 nM of non-targeting siRNA, or specific siRNAs. Transfected cells were infected/treated for the indicated time periods before being processed for analysis.

##### Wound healing model

Wounds were created in the dorsal skin of BALB/c mice using previously described protocols. On day zero, mice were anaesthetized intravenously with ketamine hydrochloride (Aneket; Neon Laboratories, Mumbai, India). Following depilation, the dorsal skin was lifted at the midline and punched through with a sterile disposable biopsy punch (4 mm in diameter; Integra Miltex), leaving one wound on either side of the midline. The wounds were then treated locally with either PBS (on the left wound) or *C. albicans* (10^5^) resuspended in PBS (on the right wound). Thus, day 0 marked the day of injury, followed by infection and subsequent observations every 24 hours. On the designated days, each wound site was digitally photographed, and wound areas were identified using ImageJ software (version 6.0). Each group (for day-to-day evaluation) included at least five mice. Wound area changes over time were calculated as a percentage of the initial wound area. Wound samples were also collected with a 6mm biopsy punch tool and crushed with RIPA buffer before being isolated and analysed by western blot. For q-RT-PCR, samples were crushed, and total RNA was isolated using TRI reagent (T9424, Sigma-Aldrich, USA). For p-CREB inhibitor topical treatment to wounds, 30μM of working concentration of the inhibitor was prepared in a mixture of DMSO and PBS (3:1). This mixture was applied to both infected and uninfected wounds on every alternate day post wounding for five days, post which images were taken, and samples were isolated for H&E staining.

##### Hematoxylin and Eosin staining

The Leica RM2245 microtome was used to cut 5 μm sections of formalin-fixed, paraffin-embedded mouse wound tissue samples. Sections were deparaffinized and rehydrated, then stained with Hematoxylin and Eosin according to the manufacturer’s instructions. Sections were dehydrated and mounted with DPX mountant. Sections were dried overnight before being handed over to a consultant pathologist for blinded analysis.

##### In vitro CFU analysis

Macrophages were given infection with *C. albicans* for zero hour (30m post infection for proper internalization) and 12h time points. Post infection, they were washed with ice cold PBS after removal of media and then lysis was done with 0.1% Triton X-100. The lysate was collected and then plated on YPD agar plate after making serial dilutions in PBS and colonies were counted for assessment.

##### Statistical analysis

Student’s t-test distribution and one-way ANOVA were used to evaluate levels of significance for sample comparisons, which were then followed by Tukey’s multiple comparisons. Data in the graphs are expressed as the mean ± S.E for values from at least three independent biological replicates, and P values < 0.05 were considered significant. All statistical analyses were performed using the GraphPad Prism 10.0 program.

## Author Contributions

A.G. and K.N.B. conceived and designed the experiments. A.G. and A.S. performed experiments. A.G., A.S. and K.N.B. wrote the manuscript and K.N.B. supervised the study.

## Acknowledgments

We thank Central Animal Facility of IISc for providing and maintaining mice for our experimentation.

## Abbreviations

KDM6A: Lysine Demethylase 6A

UTX: Ubiquitously Transcribed tetratricopeptide repeat on chromosome X

CREB: c-AMP Response Element Binding Protein

MMPs: Matrix Metallo Proteases

TIMPs: tissue inhibitors of metalloproteases

## References

1. Cuzzell, J. Infection in chronic wounds: Controversies in diagnosis and treatment. Dermatol Nurs 11, 382–382 (1999).

2. Wei, Y. et al. The clinical effectiveness and safety of using epidermal growth factor, fibroblast growth factor and granulocyte-macrophage colony stimulating factor as therapeutics in acute skin wound healing: a systematic review and meta-analysis. Burns Trauma 10, 1980–2019 (2022).

3. Cabral-Pacheco, G. A. et al. The Roles of Matrix Metalloproteinases and Their Inhibitors in Human Diseases. Int J Mol Sci 21, 1–53 (2020).

4. Schultz, G. S., Chin, G. A., Moldawer, L. & Diegelmann, R. F. Principles of Wound Healing. Diabetic Foot Problems 395–402 (2011) doi:10.1142/9789812791535_0028.

5. Barrientos, S., Stojadinovic, O., Golinko, M. S., Brem, H. & Tomic-Canic, M. Growth factors and cytokines in wound healing. Wound Repair Regen 16, 585–601 (2008).

6. Seidel, D. et al. Impact of climate change and natural disasters on fungal infections. Lancet Microbe doi:10.1016/S2666-5247(24)00039-9.

7. Dowd, S. E. et al. Survey of fungi and yeast in polymicrobial infections in chronic wounds. J Wound Care 20, 40–47 (2011).

8. Schaller, M., Borelli, C., Korting, H. C. & Hube, B. Hydrolytic enzymes as virulence factors of Candida albicans. Mycoses 48, 365–377 (2005).

9. Soll, D. R. The role of phenotypic switching in the basic biology and pathogenesis of Candida albicans. J Oral Microbiol 6, 1–12 (2014).

10. Pericolini, E. et al. Secretory Aspartyl Proteinases Cause Vaginitis and Can Mediate Vaginitis Caused by Candida albicans in Mice. mBio 6, (2015).

11. Desai, J. V., Mitchell, A. P. & Andes, D. R. Fungal biofilms, drug resistance, and recurrent infection. Cold Spring Harb Perspect Med 4, (2014).

12. Mahadik, K., Yadav, P., Bhatt, B., Shah, R. A. & Balaji, K. N. Deregulated AUF1 Assists BMP-EZH2-Mediated Delayed Wound Healing during Candida albicans Infection. J Immunol 201, 3617–3629 (2018).

13. Larionova, I., Kazakova, E., Patysheva, M. & Kzhyshkowska, J. Transcriptional, Epigenetic and Metabolic Programming of Tumor-Associated Macrophages. Cancers (Basel) 12, 1–40 (2020).

14. Rando, O. J. & Verstrepen, K. J. Timescales of genetic and epigenetic inheritance. Cell 128, 655–668 (2007).

15. Shaw, T. & Martin, P. Epigenetic reprogramming during wound healing: loss of polycomb-mediated silencing may enable upregulation of repair genes. EMBO Rep 10, 881–886 (2009).

16. Boudra, R. & Ramsey, M. R. Focus: Skin: Understanding Transcriptional Networks Regulating Initiation of Cutaneous Wound Healing. Yale J Biol Med 93, 161 (2020).

17. Wen, A. Y., Sakamoto, K. M. & Miller, L. S. The Role of the Transcription Factor CREB in Immune Function. J Immunol 185, 6413 (2010).

18. Brubaker, A. L., Carter, S. R. & Kovacs, E. J. Experimental Approaches to Tissue Injury and Repair in Advanced Age. Methods Mol Biol 1343, 35 (2015).

19. Schilrreff, P. & Alexiev, U. Chronic Inflammation in Non-Healing Skin Wounds and Promising Natural Bioactive Compounds Treatment. Int J Mol Sci 23, 4928 (2022).

20. Krzyszczyk, P., Schloss, R., Palmer, A. & Berthiaume, F. The Role of Macrophages in Acute and Chronic Wound Healing and Interventions to Promote Pro-wound Healing Phenotypes. Front Physiol 9, (2018).

21. Caley, M. P., Martins, V. L. C. & O’Toole, E. A. Metalloproteinases and Wound Healing. Adv Wound Care (New Rochelle) 4, 225 (2015).

22. Choi, I. Y. et al. Activation of cannabinoid CB2 receptor-mediated AMPK/CREB pathway reduces cerebral ischemic injury. Am J Pathol 182, 928–939 (2013).

23. Turcotte, C., Blanchet, M. R., Laviolette, M. & Flamand, N. The CB2 receptor and its role as a regulator of inflammation. Cellular and Molecular Life Sciences 73, 4449 (2016).

24. Turcotte, C., Blanchet, M. R., Laviolette, M. & Flamand, N. The CB2 receptor and its role as a regulator of inflammation. Cell Mol Life Sci 73, 4449 (2016).

25. Jozic, I., et al. Glucocorticoid-mediated induction of caveolin-1 disrupts cytoskeletal organization, inhibits cell migration and re-epithelialization of non-healing wounds. Commun Biol 4, (2021).

26. Agger, K. et al. UTX and JMJD3 are histone H3K27 demethylases involved in HOX gene regulation and development. Nature 449, 731–734 (2007).

27. Ezponda, T. et al. UTX/KDM6A Loss Enhances the Malignant Phenotype of Multiple Myeloma and Sensitizes Cells to EZH2 inhibition. Cell Rep 21, 628–640 (2017).

28. Chen, J. et al. Kdm6a suppresses the alternative activation of macrophages and impairs energy expenditure in obesity. Cell Death Differ 28, 1688–1704 (2021).

29. Schultz, G. S. et al. Wound bed preparation: a systematic approach to wound management. Wound Repair Regen 11 **Suppl 1**, (2003).

30. Raziyeva, K. et al. Immunology of Acute and Chronic Wound Healing. Biomolecules 11, (2021).

31. Guo, S. & DiPietro, L. A. Factors affecting wound healing. J Dent Res 89, 219–229 (2010).

32. Gardner, S. E., Frantz, R. A. & Doebbeling, B. N. The validity of the clinical signs and symptoms used to identify localized chronic wound infection. Wound Repair Regen 9, 178–186 (2001).

33. Koga, Y. et al. CREB regulates TNF-α-induced GM-CSF secretion via p38 MAPK in human lung fibroblasts. Allergology International 65, 406–413 (2016).

34. Bannister, S., Messina, N. L., Novakovic, B. & Curtis, N. The emerging role of epigenetics in the immune response to vaccination and infection: a systematic review. Epigenetics 15, 555– 593 (2020).

35. Bannister, A. J. & Kouzarides, T. Regulation of chromatin by histone modifications. Cell Research 2011 21:3 21, 381–395 (2011).

36. Esteller, M. Cancer epigenomics: DNA methylomes and histone-modification maps. Nat Rev Genet 8, 286–298 (2007).

